# Dual process coding of recalled locations in human oscillatory brain activity

**DOI:** 10.1101/361774

**Authors:** Mary H. MacLean, Tom Bullock, Barry Giesbrecht

## Abstract

A mental representation of the location of an object can be constructed using sensory information selected from the environment and information stored internally. Human electrophysiological evidence indicates that behaviorally relevant locations, regardless of the source of sensory information, are represented in alpha-band oscillations suggesting a shared process. Here we present evidence from human subjects of either sex for two distinct alpha-band based processes that separately support the representation of location, exploiting sensory evidence sampled either externally or internally.

**Significance Statement:** Our sensory environment and our internal trains of thought are coded in patterns of brain activity and are used to guide coherent behavior. Oscillations in the alpha frequency band are a predominant feature of human brain activity. This oscillation plays a central role in both selective attention and working memory, suggesting that these important cognitive functions are mediated by a unitary mechanism. We show that the alpha oscillation reflects two distinct processes, one that is supported by continuous sampling of the external sensory environment, and one that is based on sampling from internal representations coded in visual short-term memory. This represents a significant change in our understanding of the nature of alpha oscillations and their relationship to attention and memory.

## Introduction

Mental representations of behaviorally relevant visual features and locations are based on information sampled from the environment and can persist in visual working memory even in the absence of maintained external sensory input (Foster, Sutterer, Serences, Vogel, & Awh, 2016; Harrison & Tong, 2009; Serences, Ester, Vogel, & Awh, 2009). Unlike features (e.g. colors), when stimulus locations are being maintained in working memory, the location of a stimulus remains available so long as there is visual input from the environment to be sampled, even if the stimulus is not present. Importantly, uninterrupted visual input is not necessary for successful recall of location, as multiple attended locations can be recalled when eyes are closed during retention (Pearson & Sahraie, 2003). Thus, mental representations of locations can be constructed from information sampled from the external environment or from information encoded internally.

Both the selection (i.e., sampling) and retention of mental representations of locations, regardless of the source of information (i.e., external or internal), appear to be supported by processes associated with alpha band oscillations (Foster et al., 2016; Foster, Sutterer, Serences, Vogel, & Awh, 2017; Kelly, Lalor, Reilly, & Foxe, 2006; Samaha, Sprague, & Postle, 2016; Thut, Nietzel, Brandt, & Pascual-Leone, 2006; van Moorselaar et al., 2018). The retention of spatially selective representations, and the associated alpha activity, may represent a continuous process of selection, i.e., the focus of attention (Lewis-Peacock, Drysdale, Oberauer, & Postle, 2012), working memory, or a combination of both. Consistent with this hypothesis, recent evidence from a dual task paradigm involving a shift of spatial attention during the retention interval of a spatial working memory task revealed that alpha carried information about the location stored in memory, but then when attention was cued to a new location, alpha carried information about the newly attended location until the task was complete and shifted back to the location stored in memory (van Moorselaar et al., 2018). This systematic tradeoff between the attention and memory tasks suggests that alpha represents a unitary mechanism that mediates spatial selectivity and that this mechanism is shared by attention and spatial working memory.

While the evidence reviewed above is consistent with the notion that alpha represents a unitary mechanism for the selection and retention of behaviorally relevant locations, this evidence comes exclusively from tasks in which there is sustained visual input from the external environment. Thus, it is unknown whether similar processes are involved in coding information from internal representations, as may be required when sensory input is disrupted (e.g., by eye closure or sensory degradation) and, as such, it is unclear whether alpha represents a unitary process for the selection and retention of locations when information is sampled from the external environment or from information that has already been encoded internally (i.e., information in visual working memory). Here we present evidence that at least two distinct processes associated with alpha-band oscillations support spatially selective representations: (1) a fast and continuous location selection process that is disrupted by a masking stimulus and that degrades without continued external visual input, and (2) a delayed process that emerges when external visual input is no longer available. Participants performed a delayed spatial estimation task (Foster et al., 2016; Wilken & Ma, 2004; Zhang & Luck, 2008), in which a single stimulus was briefly presented in the periphery, in one of eight possible locations on the circumference of an imaginary circle (**Fig. 1a**). Participants were instructed to attend covertly to the stimulus location, remember its location, and report the location after a short delay. The presence or absence of a masking checkerboard after the stimulus, and the state of the eyes during retention (open or closed) yielded four within-subject conditions: eyes open, eyes open + mask, eyes closed, eyes closed + mask.

**Figure 1.**
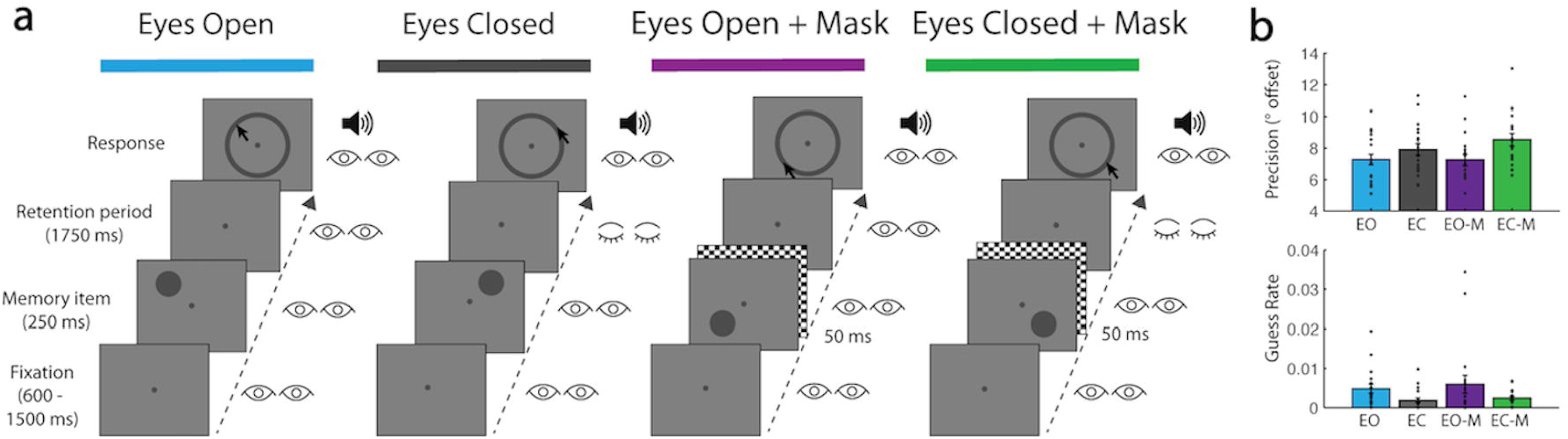
Delayed spatial estimation task. **(a)** Trial procedure for each experimental condition. **(b)** Precision of spatial estimation and guess rate. Error bars = standard error of the mean.

## Materials & Methods

### Participants

Participants were 18 adult volunteers (mean age = 20.8 years, 13 females, all right handed) recruited from the University of California Santa Barbara community. Participants were compensated at a rate of $10 per hour (minimum of $120 total/participant). All participants reported normal or corrected-to-normal vision. The UCSB Human Subjects Committee approved all procedures.

### Stimuli and procedure

Each participant completed four testing sessions on four separate days. Sequential sessions were separated by a minimum of 24 hours and a maximum of 1 week. All four sessions were completed within an eight-week period for a given participant. In each session the participant was required to perform 15 blocks of 64 trials (960 trials total) of the same delayed spatial estimation task. Each session consisted of a different experimental condition of the task: “eyes open” (EO), “eyes closed” (EC), “eyes open with mask” (EO+M), “eyes closed with mask” (EC+M). The order of experimental conditions was counterbalanced between participants.

Each trial of the delayed spatial estimation task began with a small blue fixation dot (subtending 0.2° visual angle) in the center of the screen, along with a green dot (subtending 0.4° visual angle) representing the location of the participant’s gaze. The participant aligned their gaze dot with the fixation dot and pressed the space bar to start the trial. The fixation dot immediately turned grey to indicate that the trial was underway. After a brief fixation (jittered randomly between 0.6 s and 1.5 s) the stimulus, a grey target circle (subtending 1.6° visual angle), appeared, centered at a point on an imaginary circle circumventing 4° from fixation. The stimulus appeared within one of eight angular location bins relative to fixation [0°, 45°, 90°, 135°, 180°, 225°, 270°, 315°], with angle jittered randomly within each bin between +1-44°.

The stimulus remained on screen for 0.25 s. Following stimulus offset there was a 1.75 s retention interval. A grey response ring then appeared, circumventing 4° of visual angle from fixation, along with an auditory cue (0.1 s, 250 Hz pure sine-wave). The participant used the mouse cursor to indicate the location of the target on the response ring. The mouse cursor appeared in a randomized corner of the response screen on each trial. This prevented the participant from anticipating the forthcoming mouse movement during the retention period. The key manipulations between sessions occurred immediately post-stimulus, during the retention interval. In the EO condition participants were instructed to maintain fixation for the duration of the retention interval. In the EC condition participants were instructed to close their eyes immediately after stimulus offset and reopen them when they heard an auditory cue at the end of the retention interval. In the EO+M and EC+M conditions participants were given then same respective instructions, however the target was masked with a global checkerboard (0.05 s, whole screen, each checker subtending 0.48° by 0.48° of visual angle).

Gaze-contingent eye tracking was employed in all sessions during fixation, stimulus presentation and retention periods. Trials where participants blinked or their gaze deviated more than 2.4° from fixation were aborted. In the EC/EC+M conditions, the trial was also aborted if the participant did not close their eyes between 0.05 s and 0.5 s following target offset. This ensured that participants closed their eyes in a timely manner and also that they viewed the checkerboard mask in its entirety in the masked condition. If the participant reopened their eyes during the retention period of the EC/EC+M condition the trial was aborted. On all trials where the gaze-contingent conditions were broken, the next trial began immediately and the aborted trial was added to the end of the trial sequence for that block of trials, thus ensuring a complete set of trials without gaze deviations were recorded for each participant in each condition.

While performing the task the participant was positioned in a chin rest at 120 cm viewing distance from a monitor (19 inch ViewSonic E90f CRT), and stimuli were presented using custom scripts that utilized the Psychophysics Toolbox for MATLAB. An eye-tracker (Eyelink 1000 plus, SR Research Ltd, Mississauga, Ontario, Canada) was positioned 60 cm from the right eye (monocular tracking @ 1000 Hz, mean error < 1°). Performance on the delayed spatial estimation task was modeled using the MemToolbox (Foster et al., 2016; Suchow, Brady, Fougnie, & Alvarez, 2013; Zhang & Luck, 2008). A standard mixture model with bias was applied to error in the spatial estimates (in degrees of offset) of each participant within each condition yielding a measure of precision of estimation as well as guess rate.

### EEG Data Acquisition

EEG data were recorded for each participant using a BioSemi Active Two system (BioSemi, Amsterdam, Netherlands) consisting of 64 Ag-AgCl sintered active electrodes arranged in an elastic cap (Electro-Cap, USA) and placed in accordance to the 10-20 system. Additional electrodes were placed at the right and left mastoids, as well as 1 cm lateral to the left and right canthi (horizontal) and above and below each eye (vertical) for the EOG. Data were sampled at 1024 Hz and referenced offline to the average mastoid signal. At the beginning of each investigation all impedances were < 20 kΩ. All recording took place in an electrically shielded chamber to ensure minimal interference from external sources of electrical noise.

### EEG Data Pre-processing

Custom scripts in MATLAB (version 2013b, Massachusetts, The MathWorks Inc.) and using the EEGLAB toolbox (Delorme & Makeig, 2004) were used for offline processing of the EEG data. The continuous data were referenced to the average mastoid signal and then high- and low-pass filtered between 0.1 Hz and 80 Hz, respectively (EEGLAB function *pop_eegfiltnew)*. The Automatic Artifact Removal (AAR) toolbox (Gomez-Herrero et al., 2006) was then used to remove ocular artifacts associated with eye closure during the EC/EC-M conditions. The data were then resampled at 256 Hz (EEGLAB function *pop_resample*), to reduce computation time and memory demands, then epoched between −0.5 and 2.5 s around the onset of the stimulus and (EEG data for aborted trials were discarded). The data were then visually inspected and noisy electrodes were excluded (mean electrodes excluded [mean ± SEM] = 1.23 ± .36). In order to maintain consistency in the topographic distribution of electrical activity across the scalp, noisy electrodes that were excluded from one condition were also excluded from all other conditions for that subject. Trials exceeding ± 150 µV in remaining electrodes were then excluded (mean trials excluded: 19.21 ± 4.53; EC 17.06 ± 3.15, EO+M 16.27 ± 3.27; EC+M 16.06 ± 4.14).

### Inter-Trial Coherence

In order to observe the evoked response to the stimulus, the mask, and eye closure we calculated inter-trial coherence (ITC) of the epoched data in the alpha (8-12 Hz) range (EEGLAB function *erpimage*) for each participant within each condition. ITC reflects the phase coherence of the EEG signal between trials (between 0 and 1, where 0 indicates no coherence and 1 indicates perfect coherence). Increased phase coherence reflects a reset in phase time-locked to an event (stimulus, mask or eye closure onset). This value is determined separately within each electrode. As we were interested in signals evoked by visual processing we averaged the ITC in parietal and occipital electrodes (P07/8, P03/4, O1/2, POz, Oz).

### Spectral Analysis

Epoched data were filtered using a 3rd order Butterworth bandpass filter (MATLAB function *butter*) from 8-12 Hz (alpha). A Hilbert transformation (MATLAB function *hilbert*) was then applied to the filtered signal in order to obtain a measure of instantaneous amplitude and phase. The data were then epoched again between −0.5 to 2 s (.5 s pre stimulus to the end of the retention period).

Prior to modeling, induced power was calculated as the square of the absolute of the Hilbert transformed complex values. Induced power reflects ongoing oscillatory activity regardless of its phase relationship with stimulus onset. Within each location bin trials were then randomly subdivided into three samples and then averaged to create three averaged trials per location bin. Following averaging, each condition included 24 averaged trials (8 location bins x 3 averaged trials).

Because trial-based artifact rejection results in uneven numbers of trials per condition, it was necessary to ensure that any comparisons between conditions were not influenced by unequal trial counts. Prior to entering the data into the IEM, the minimum number of trials per location bin (*n)* was calculated across the four conditions for each participant. To ensure equal numbers of trials from each location bin were entered into the model, *n*-1 trials were randomly selected from each bin.

### Inverted Encoding Model

An IEM was used to estimate spatially selective neural population (“channels”) response profiles based on the topographical distribution of alpha power across the scalp (Foster et al., 2016). The model first estimates the extent to which the linear combination of a priori canonical channel responses (i.e. a basis set) captures the underlying structure of the observed data (topographical distribution of alpha power), yielding a set of regression weights. The model then uses these weights to estimate the channel response from the observed data. The parameters of these channel response estimates can then be used to quantify the spatially selective response. This method was initially applied to fMRI data (Brouwer & Heeger, 2009, 2011, 2013; Ester, Sprague, & Serences, 2015; Naselaris, Kay, Nishimoto, & Gallant, 2011; Serences & Saproo, 2012) and has been recently applied to EEG recorded at the scalp (Bullock, Elliott, Serences, & Giesbrecht, 2017; Foster et al., 2016, 2017; Garcia, Srinivasan, & Serences, 2013; Samaha et al., 2016).

The IEM was run separately for each sample in time (256 Hz EEG sampling rate x 2.5 sec = 640 samples) using induced alpha power within each condition for each participant. Evoked power was also modeled; the results replicated those of Foster et al. (2016), and did not differ by condition. Note that as performance of the IEM is sensitive to noise we compared the alpha signal-to-noise ratio (SNR; alpha power / standard deviation of power in other frequencies) between conditions and found no difference (all effects BF < 1). For cross-validation of the IEM a *k*-fold approach was employed where *k* = 3. The averaged trials were randomly grouped into three blocks, each with one averaged trial per location bin. Training was performed using 2/3 blocks and the resulting model was tested on the remaining block. This was repeated such that each block served as the test block.

For each participant and condition, *m* represents the number of EEG electrodes in each dataset (mean across participants = 62.77 ± .36; equivalent within participant), *n_1_* represents the number of trials in the training set (2 blocks of 8 trials) and *n_2_* represents the number of trials in the testing set (1 block of 8 trials). Let *j* be the number of hypothetical location selective channels (C_1_, *j* x *n_1_*), composed of half-sinusoidal functions raised to the seventh power as the basis set. Here, the basis set was comprised of 8 equally spaced locations (i.e. *j*=8). B_1_ (*m* x *n_1_*) represents the training set and B_2_ (*m* x *n_2_*) the test set. A standard implementation of the general linear model (GLM) was then used to estimate the weight matrix (W, *m* x *j*) using the basis set (C_1_). More specifically, using the GLM:

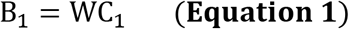

Then, the ordinary least-squares estimate of W can be computed as:

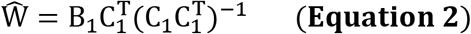

Using the estimated weight matrix (Ŵ, **Equation 2**) and the test data (B_2_), the channel responses C_2_ (*j* x *n_2_*) can be estimated by:

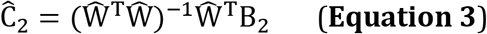

After the Ĉ_2_ was solved for each location bin, the channel response function on each averaged trial was then circularly shifted to a common stimulus-centered reference frame (degrees of offset from channel’s preferred location bin), and the centered response functions were averaged across channels. The model was repeated for each time sample. To safeguard that the outcome of the model and ensure that any subsequent analyses were not influenced by an idiosyncratic selection of trials, this process was repeated 10 times, with a randomized selection of trials entered into the IEM for each of the iterations, and the final centered tuning function was computed by averaging over the 10 iterations.

The IEM procedure was also performed with randomly permuted location bin labels. Randomizing the location bin labels should eliminate spatially selective responses and result in flat channel response profiles. These permuted responses were used in hypothesis testing.

### Temporal Generalization of IEM

In order to observe the intra- and inter-condition temporal generalization of the IEM we ran the IEM as described above by training at each sample in time, but then testing on every sample in time, either within condition or between conditions. In order to reduce computation time and the number of comparisons we down-sampled the data prior to modeling from 256 Hz (640 samples) to 16 Hz (40 samples). We repeated the generalization IEM procedure with permuted location bin labels for the purpose of hypothesis testing.

### Quantifying Spatially Selective Representations

To quantify spatial selectivity the estimated channel responses were folded around 0° of offset, transforming the responses from [−135, −90, −45, 0, 45, 90, 135, 1. 180] into [0, 45, 90, 135, 180] by averaging the response at corresponding offsets (± 45, 90, and 135°; 0° an 180° were not averaged). Slope was then computed (MATLAB function *polyfit*) as the linear regression weight of alpha power across offset. Larger slope values indicate greater spatial selectivity.

### Linear Decoding Analysis

To assess the extent to which the scalp topography of alpha power discriminated between the neural responses to stimuli presented in the eight different location bins without assuming an a priori response profile (i.e., the basis set in the IEM approach), we trained and tested linear discriminant pattern classifiers in a series of three analyses that paralleled the IEM analyses. First, we trained and tested classifiers within each condition at each point in time during the trial. Second, we conducted an intra-condition temporal generalization analysis in which a classifier was trained at each time point and tested on all of the other time points. Third, we conducted an inter-condition temporal generalization analysis in which training was conducted on each time point in one condition and tested on all the time points in each of the other conditions.

For each analysis, the classifier (MATLAB Statistics and Machine Learning Toolbox *classify, ‘linear’ option)* was trained using power at parietal (P1-10,z), parietal-occipital (PO3,4,7,8,z), occipital (O1,2,z), and inion (Iz) electrodes. We restricted the analyses to these 20 electrodes on an a priori basis to reduce the dimensionality of the training data, ensuring stability of the classifier. To reduce computation time and to facilitate comparison with the IEM generalization analyses we down-sampled the data at 16 Hz, (16 Hz EEG sampling rate x 2.5 sec for each trial = 40 samples). Leave-one-trial-out cross validation was used to train and test the classifiers and performance was measured by computing the proportion of correct classifications (n correct classifications/total classifications). Chance for these analyses was 1/8=0.125. As with the IEM analyses, all classification analyses were rerun permuting the location bin labels.

### Alpha Lateralization Analysis

Systematic changes in alpha power topography that occurred as a function of stimulus locations were assessed by normalizing (i.e., dividing) the difference in alpha power at contralateral and ipsilateral parietal/occipital electrodes sites (PO3/4, PO7/8, O1/2) by the sum of power at contralateral and ipsilateral sites. Normalized alpha power at contralateral and ipsilateral sites was then averaged by condition and time window. The first time window overlapped with cue presentation, following the earliest stimulus-related evoked potentials and the emergence of a spatially selective representation (198-250 ms). The second time window overlapped with the retention interval (post-mask/eye closure; 750-2,000 ms). Because the first time window is only 52 ms, while the second is 1,250 ms, a 52 ms window was randomly and independently selected from the retention interval for each condition for each participant, such that the average latency of the window was centered ∼ 1,375 ms after stimulus onset. This ensured that estimates of the mean and variability in each time window were based on the same number of time points.

### Experimental Design and Statistical Analysis

Bayes Factor (BF) was calculated for the purpose of inferential statistics (Kass & Raftery, 1995; Kruschke & Liddell, 2016; Rouder, Morey, Speckman, & Province, 2012) using functions from the BayesFactor toolbox for R(Morey, Rouder, & Jamil, 2015), which employs a Cauchy prior. According to various guidelines a BF between 1-3 indicates “anecdotal evidence” for H1, between 3 and 10 indicates “moderate” evidence, between 10 and 30 indicates “strong” evidence and greater than 30 indicates “very strong” evidence (Dienes, 2016; Kass & Raftery, 1995; Kruschke & Liddell, 2016; Wetzels et al., 2011).

To test the reliability of the spatially selective channel responses, BF paired t-tests (R function *ttestBF*) were computed at each sample in time comparing the slopes of the real and permuted channel response estimates. To test for differences in slope between conditions BF ANOVAs (R function *anovaBF*) with the factors eyes (open vs. closed) and mask (masked vs. unmasked) were computed comparing the slopes of the real channel response estimates at each time point. To contrast slope values resulting from the temporal generalization analyses within conditions, to those between conditions, BF paired t-tests were also computed for each cell in the generalization matrix. To assess the relationship between slope and ITC, BF regressions (R function *regressionBF*) were computed. BF ANOVAs and t-tests were also used to test for differences in performance.

Our analyses involve repeated comparisons at multiple time points, which raises the specter of increased inferential error. However, it should be noted that Bayesian inference, even without correcting for multiple comparisons, is more conservative than frequentist inference, and much less likely to result in false confidence (Gelman & Tuerlinckx, 2000). Furthermore, our effects are present at multiple time points, as hypothesized, and no inference relies on a single comparison, but rather a pattern of effects across time. Our inferences are further strengthened by our use of IEM parameters based on permuted data for comparisons, which yields a more realistic null for our comparisons, and that our parameters are averaged over multiple repetitions of the IEM. These features should help mitigate errors due the peculiarities in the data at a given time point or sample of trials.

### Code Accessibility

All custom code will be made available in a public repository on GitHub/OSF, please contact the corresponding author for more information.

### Control Study

Participants in the control study were 12 adult volunteers (mean age = 22.9; six males; 11 right-handed) recruited from the University of California Santa Barbara community. Participants were compensated at a rate of $10 per hour (minimum of $120 total/participant). All participants reported normal or corrected-to-normal vision. The UCSB Human Subjects Committee approved all procedures.

Participants completed two conditions of the same delayed spatial estimation task as described above, in two separate sessions on two different days: (1) eyes open, and (2) eye-blink. The eyes open condition was identical to that used in the main experiment. The eye-blink condition is identical to the eyes closed condition except that participants were instructed to re-open their eyes after closing them in response to the auditory cue (i.e., we asked them to blink). The deadline for re-opening their eyes was the same as the deadline for eye closure in the eyes closed conditions (750 ms post cue). Performance was unaffected by condition (precision: Eyes Open = 7.16 ± .41, Eyes Blinked = 6.66 ± .40, BF_t-test_ = 1.61; guess rate: Eyes Open = .004 ± .001, Eyes Blinked = .002 ± .002, BF _t-test_ < 1). All EEG recording and analyses, as well as the approach to statistical inference, were identical to those described above.

## Results

### Location report precision and guess rate are affected by eye closure

The precision of the reported stimulus location was reduced when participants were required to close their eyes (*BF* > 1,000), an effect enhanced by the presence of a mask (*BF* > 1,000; **Fig. 1b**). This suggests that the representation of location without the support of continued external visual input is less precise, especially when masked. While guess rate was extremely low in all conditions, there was a small decrease in guessing when participants were required to close their eyes during retention (BF > 7). Modulation of recall precision by eye closure is inconsistent with evidence that spatial recall, when measured as accuracy, is robust to eye closure without saccades (Pearson & Sahraie, 2003); however a continuous report measure, as in the case of the delayed estimation task used here, is possibly more sensitive to the effects of eye closure than discrete measures.

### Reconstructions of location representations are temporarily disrupted by a visual mask, and degrade over time with eye closure

In order to observe the spatially selective representation of the stimulus location, an inverted-encoding model (IEM) was employed to reconstruct spatially selective responses from topographically specific patterns of alpha band oscillations in scalp-recorded EEG (Foster et al., 2016; Samaha et al., 2016; van Moorselaar et al., 2018) **(Fig. 2a-c**). More specifically, the IEM method used here employed a general linear model in which the topographic pattern of oscillatory activity was regressed onto an *a priori* defined set of spatially selective channels. This basis set consists of the expected response of each channel to each location as expressed in arbitrary units – i.e., the hypothesized response function of each spatially selective channel. The shape of the hypothesized function resembles that of a tuning function as observed in neural population responses, where the channel is activated the most in response to its unique preferred location with decreasing activation at increasingly distant locations, and is recreated as a half sine function raised to some power. The resulting regression weights, trained and tested using leave-one-out cross-validation, reflect the pattern of neural activity particular to each spatially selective channel, i.e., only that variability predicted by the basis set. The product of the weights and the observed neural activity yields an estimate of the response in each spatially selective channel – i.e., the representation of location.

**Figure 2.**
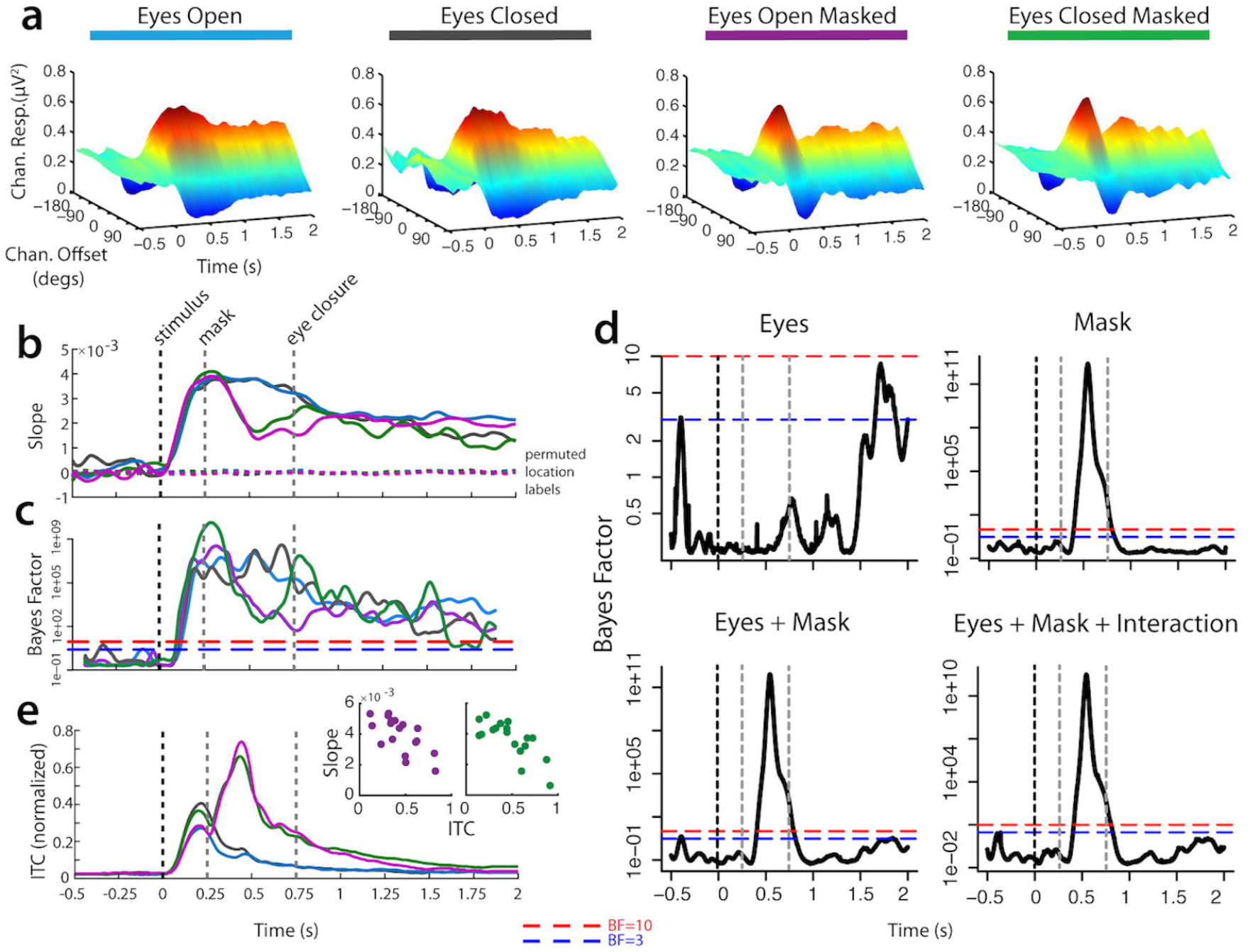
Results of the alpha-based inverted encoding model estimation of the stimulus location. **(a)** Estimated neural population response, or “channel” (offset of channel’s preferred location from stimulus location: ±0,45,90,135, or 180°), response (alpha-band power) starting 500 ms before onset of stimulus until end of retention period. **(b)** Linear regression weights of estimated channel response folded around 0° offset (“slope”), indicating the degree of spatial selectivity. **(c)** Bayes Factor (BF) of t-test comparing slope at each time point to slope of response estimated using permuted location labels. **(d)** BF of ANOVA for effects of eye closure and mask on slope. **(e)** Inter-trial coherence (ITC) indicating stimulus- and mask-evoked potentials and scatterplots of correlation between ITC and slope during the period of the mask-evoked potentials for masked conditions.

There was evidence of a spatially selective response, centered on the stimulus location, in the alpha band following stimulus presentation under all conditions. The spatially selective response emerged ∼200 ms after stimulus onset, around the time when initial effects of covert shifts in spatial attention have previously been observed in evoked activity (Di Russo, Martínez, & Hillyard, 2003; Hillyard et al., 1999).

The presentation of a masking stimulus disrupted the spatially selective response beginning ∼200 ms after mask onset (see slope difference between masked and unmasked conditions during the .25-.75s time window in **Fig. 2b** and the corresponding BF for the effect of mask **Fig. 2d “mask”**). An evoked potential observed at parietal and occipital electrode sites, phase-locked to the onset of the mask, was observed to coincide with this disruption (**Fig. 2e**). The magnitude of the mask-evoked response predicted the magnitude of disruption to the spatially selective response shortly after mask onset (*r* (16) = -.66, *BF* > 10, and *r* (16) = -.76, *BF* > 300, for EC+M and EO+M respectively). However, this disruption was neither complete nor permanent. While degraded, evidence for a spatially selective response remained strong following the mask and returned to similar levels observed in the absence of a mask ∼500-700 ms after mask onset. This pattern suggests that while a mask disrupts the retention of a spatially selective representation, the representation can be recovered, even in the absence of external visual input when eyes were closed.

Unlike the effect evoked by the mask, there was no evidence that the spatially selective response was disrupted by eye closure (**Fig. 2d “eyes”**). Neither did eye closure evoke a phase-locked response (**Fig. 2e**). There was, however, moderate evidence of a small reduction of the spatially selective response later in the retention interval ∼1,000 ms after eye closure, as compared to when eyes remained open. Nonetheless evidence for the spatially selective representation remained robust throughout the retention interval when eyes were closed. Thus, the disruptive effect induced by eye closure on spatially selective representations is smaller and delayed compared to the immediate disruptive effect evoked by the mask.

Importantly, the results of the IEM demonstrate that a spatially selective representation was present in alpha under all conditions in the retention interval *to the same degree*, except for some evidence of a slight drop in selectivity at the very end of the interval when eyes were closed. This would appear to support the unitary nature of the spatially selective mechanism in alpha oscillations – that the representation is robust to changes in input. However, as the following section will demonstrate, while a spatially selective representation can be recreated from alpha similarly under both eyes open and eyes closed conditions, this representation is not encoded by the same pattern (topography and/or power) of alpha under both conditions.

### Reconstructions of location representations are supported by dual processes

The retention of a spatially selective representation, and its disruption by a masking stimulus, is evident in alpha band oscillations under conditions both with and without continued external visual input. Thus, the patterns of alpha band oscillations that encode for location are persistently robust throughout the entire stimulus presentation and retention interval regardless of condition. However, it is unclear whether a) the continuous encoding of location in alpha band oscillations is supported by a single common process or by multiple distinct processes, and b) which process or processes supporting the spatially selective response are affected by the disruption of external visual input (i.e., masking, eye closure, or both).

In order to observe the dynamics of the process, or processes, supporting spatially selective representations under these different conditions we examined the inter-temporal and inter-conditional generalization of the IEM. The generalization approach can be used to identify the extent that specific neural codes, reconstructions of spatial spatially selective responses in this case, are present across time and/or conditions (King & Dehaene, 2014). Models of neural codes trained on the topographic pattern of neural activity in specific time windows and/or conditions that can recover similarly reliable spatially selective patterns in other time windows and conditions would be those that exhibit generalization. Such a pattern would be evidence for a common process, as a spatially selective representation can be recreated from the same pattern of alpha as at another time/in another condition. In contrast, models trained on the topographic pattern of neural activity in specific time windows and/or conditions that fail generalize would be evidence that the spatially selective representation is coded by a different topographic pattern of neural activity predicted by the basis set, i.e. a different process. In the case of the IEM model, generalization indicates that the beta weights, representing the linear regression of alpha against the basis set, are different from one time/condition to another.

In the case where a lack of generalization is observed, it is important to rule out the possibility that the impaired ability of the model to reconstruct spatially selective patterns in other time windows and/or conditions is not merely due to a loss of signal. Importantly, there was evidence of a spatially selective response at all time points in all conditions beginning ∼200 ms after stimulus onset, thus any lack of generalization can not be attributed to a loss of the signal that encodes for location during these times in any of the conditions. While the ability to generalize between conditions may be impaired by changes in cap placement, impedance, etc. between sessions, this effect should be equivalent between all conditions at all time points.

When eyes were open and no mask was presented there was strong evidence of a *unitary* process supporting the spatially selective response during both stimulus presentation and retention intervals with continuous external visual input (**Fig. 3a “Train EO → Test EO”**), as there was extensive temporal generalization both forward and backward in time. This pattern is consistent with the conclusion that the process maintaining the spatially selective representation during retention when eyes were open is also the same process as during stimulus presentation (i.e., during selection), and remains static throughout the entire retention interval.

**Figure 3.**
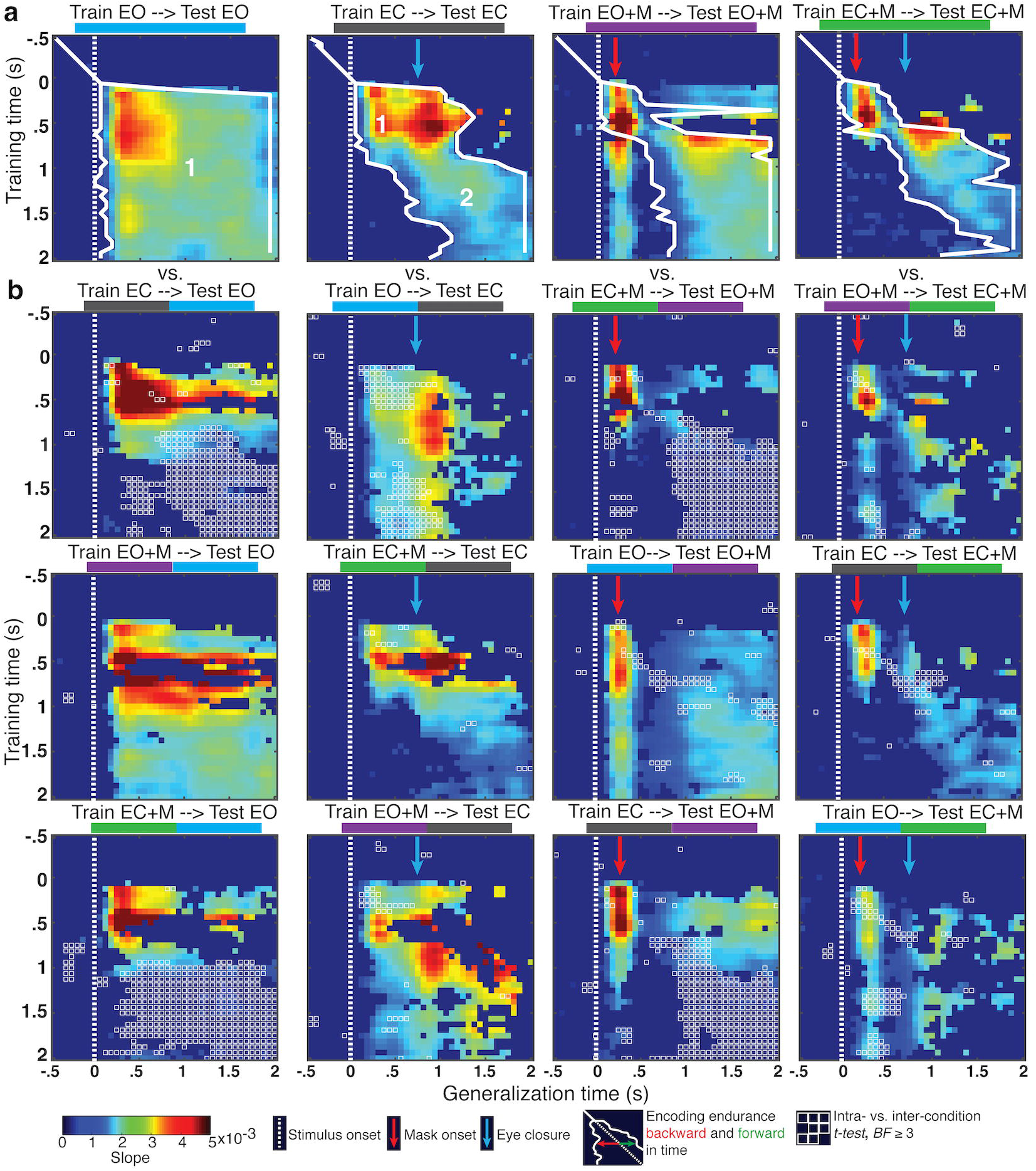
IEM generalization. **(a)** Temporal generalization within experimental condition. Slopes for a spatially selective response are plotted. Those with less than moderate evidence (BF<3) plotted as uniform dark blue. Encoding endurance indicates the sequential extent of temporal generalization backward and forward in time, where BF ≥ 3. Process 1 and 2 are indicated in the EO and EC plots. **b)** Inter-condition generalization. Points with moderate or greater evidence (BF ≥ 3) for a difference between intra-condition generalizations shown in Panel A and inter-condition generalizations are outlined in white.

Despite a similarly continuous and robust encoding of location, this was not the case without continuous visual input, when eyes were closed (**Fig. 3a “Train EC → Test EC”**). Earlier time points did not generalize forward in time to the end of the retention interval, nor did later time points generalize backward in time to the onset of the cue despite evidence for spatially selective responses at those times. Instead, when eyes were closed the pattern of temporal generalization supports two processes: a robust selection process beginning ∼100 ms after cue onset and persisting until ∼500 ms *after* eye closure before decaying; and a subsequent process, starting around the time of eye closure (∼500 ms after cue onset) and lasting the remainder of the retention interval. At this point, rather than assigning mechanistic labels to these processes (e.g., selection, retention, etc.), we take a more agnostic approach, labeling these processes in terms of their ordinal position during the trial sequence and refer to them as process 1 and process 2, respectively.

Process 1 is interrupted in the period following the presentation of a mask in both the eyes open and closed conditions (**Fig. 3a “Train EO+M → Test EO+M”/ “Train EC+M → Test EC+M”**). With eyes open, process 1 re-emerges ∼200 ms post-mask, but fails to re-emerge with eyes closed. This pattern of generalization indicates that process 1, possibly a spatial selection process, is dependent on uninterrupted external visual input, is disrupted by a visual masking stimulus, and can fully recover a spatial representation when external visual input remains available post-mask (i.e., when eyes remain open). This is similar to the observation that a subsequent cue can restore degraded representations (Sprague, Ester, & Serences, 2016), however, in the current case, there is spontaneous recovery with no aid of additional cues to location, and only when input remains.

There was strong evidence that process 1 generalized between eyes open and eyes closed conditions, regardless of mask presence (**Fig. 3b**). Process 1, trained in one eye condition and tested in another, persisted robustly forward in time towards the end of the retention interval in the eyes open conditions (**“Train EC → Test EO”/ “Train EC+M → Test EO+M”)**, but not in the eyes closed conditions (**“Train EO → Test EC”/ “Train EO+M → Test EC+M”**). Although evidence for generalization between eyes open and eyes closed conditions was stronger and more consistent in the unmasked conditions, process 1 still generalized between masked and unmasked conditions whether eyes were open or closed. This pattern suggests that process 1, the process that overlaps with stimulus processing, is common to all conditions.

Unlike process 1, there was only sporadic, weak evidence that process 2, which supported the representation of spatial selectivity during the retention period when eyes were closed, generalized to the same period when eyes were open. This may indicate that process 2 is not entirely absent in conditions with continuous external visual input, but may also be present to some degree when such input is available. However, there is evidence that process 2 generalized between the eyes closed conditions. This pattern of generalization indicates that process 2 emerges *reliably* when there is an interruption to external visual input, as when the eyes are closed.

### Evidence for dual processes in the alpha frequency band is not dependent on the IEM analytic method

The strength of the IEM approach is that the selection of the basis set is grounded in what is understood about the responses of the visual system to the feature of interest, in this case spatial location. The interpretability of the estimated channel responses is heavily dependent on the careful selection of the basis set (for a discussion of these issues please see, (Gardner & Liu, 2019; Liu, Cable, & Gardner, 2018; Sprague et al., 2018; Sprague, Boynton, & Serences, 2019). Here, we use the same basis set as that used in previous work to describe the nature of alpha power as a signal for both encoding and maintaining spatial information in memory (Foster et al., 2016, 2017; van Moorselaar et al., 2018). However, it is also important to demonstrate that the results we present here using the IEM method are not the result of any possible idiosyncrasies of that particular basis set. A common approach is to use a set of Kronecker delta functions, as the basis set which assumes that the channel responses are impulse responses, and then recreate the IEM analysis with the new basis set (Foster et al, 2016; Bullock et al., 2017). Another approach is to circumvent the IEM approach altogether and to simply determine whether the same signal (i.e., alpha power) can be used to decode the stimulus locations. The assumption in this latter approach is that the responses to the different stimulus locations are discriminable in some way that doesn’t necessarily map onto any specific a priori response profile.

We repeated the generalization analyses using a linear discriminant classifier for location, using classifier accuracy as the primary dependent variable instead of slope (**Fig. 4**). The pattern of results from the decoding analysis was very consistent with the IEM analysis. Within condition, there was strong evidence for classifier generalization across all time windows in the **EO** condition, but not in the **EC** condition (**Fig. 4a**); and there was also evidence that the early spatial selection process is disrupted by the mask and can recover later in the retention interval. The inter-condition temporal generalization decoding analysis (**Fig. 4b**) also provided evidence consistent with the IEM analysis, such that classifiers trained during the early time window generalized across conditions during those same time windows, but those trained on later time windows did not.

**Figure 4.**
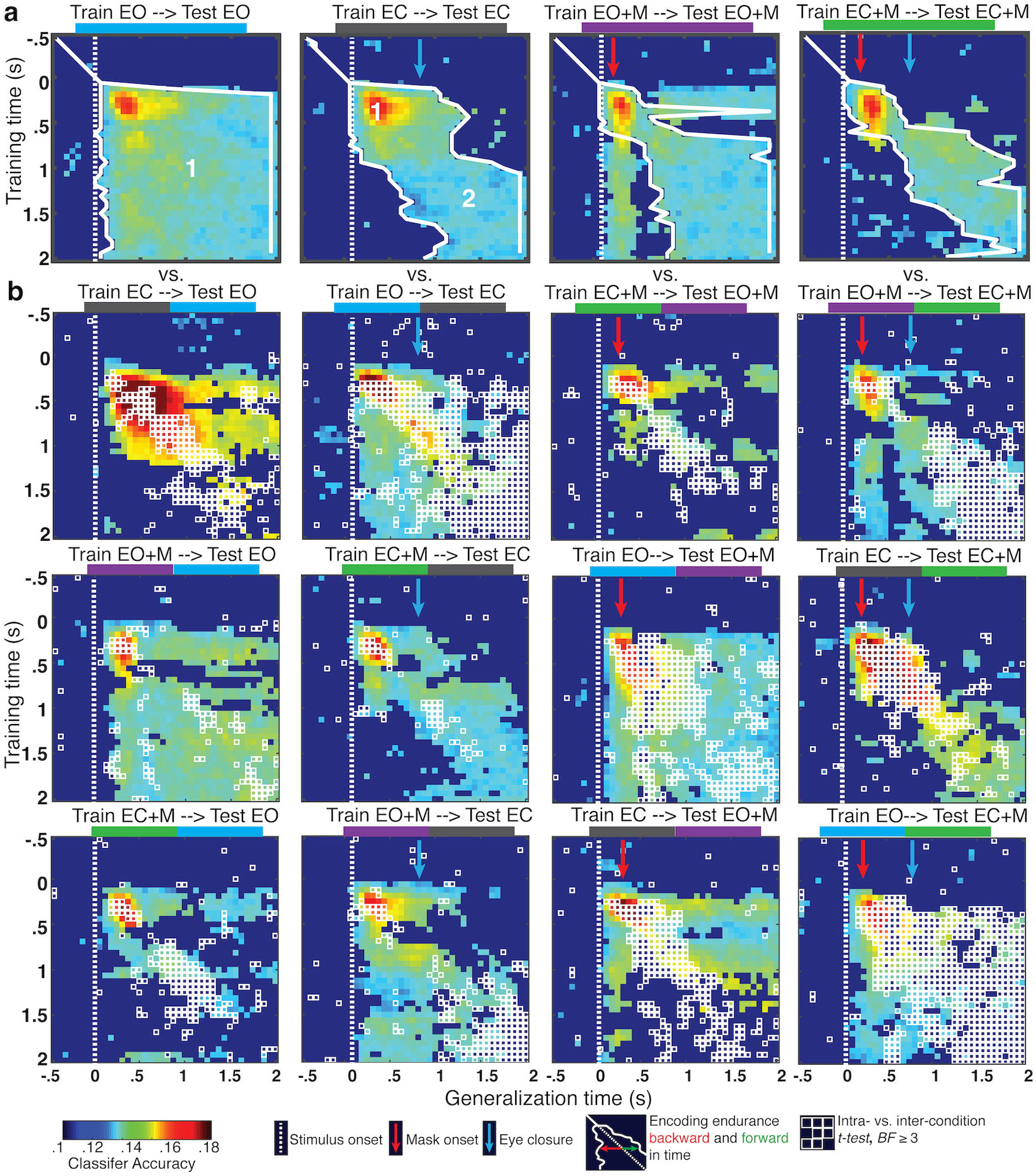
Linear discriminant classifier generalization. **(a)** Temporal generalization within experimental condition. Classifier accuracy is plotted. Accuracy with less than moderate evidence (BF<3) plotted as uniform dark blue. Encoding endurance indicates the sequential extent of temporal generalization backward and forward in time, where BF ≥ 3. Process 1 and 2 are indicated in the EO and EC plots. **b)** Inter-condition generalization. Points with moderate or greater evidence (BF ≥ 3) for a difference between intra-condition generalizations shown in Panel A and inter-condition generalizations are outlined in white.

Thus, as with the IEM analysis, the pattern of temporal generalization in the linear decoding analysis supports a series of two processes: (process 1) a robust selection process beginning ∼100 ms after cue onset and persisting until ∼500 ms *after* eye closure before decaying; and (process 2) a subsequent process, starting around the time of eye closure (∼500 ms after cue onset) and lasting the remainder of the retention interval.

### Changes in alpha, if unrelated to the code for the spatially selective representations, are insufficient to disrupt temporal generalization

Eye closure results in a well-documented increase in alpha power that is not uniformly distributed, rather it is larger at posterior electrodes relative to frontal and temporal electrodes (Barry, Clarke, Johnstone, Magee, & Rushby, 2007; Chapman, Armington, & Bragdon, 1962). Here, as expected, alpha power increased with eye closure (see **Fig. 5a**) and this increase in power was localized primarily in parietal and occipital electrodes (i.e., over visual cortex; **Fig. 5b**). Importantly, these electrodes are also associated with the largest GLM weights from the IEM (see **Fig. 8c**). Thus, it is reasonable to wonder whether such changes in alpha power and/or topography resulting from eye closure could have influenced the coding of the spatially selective representation in alpha and its ability to generalize across time and conditions.

**Figure 5.**
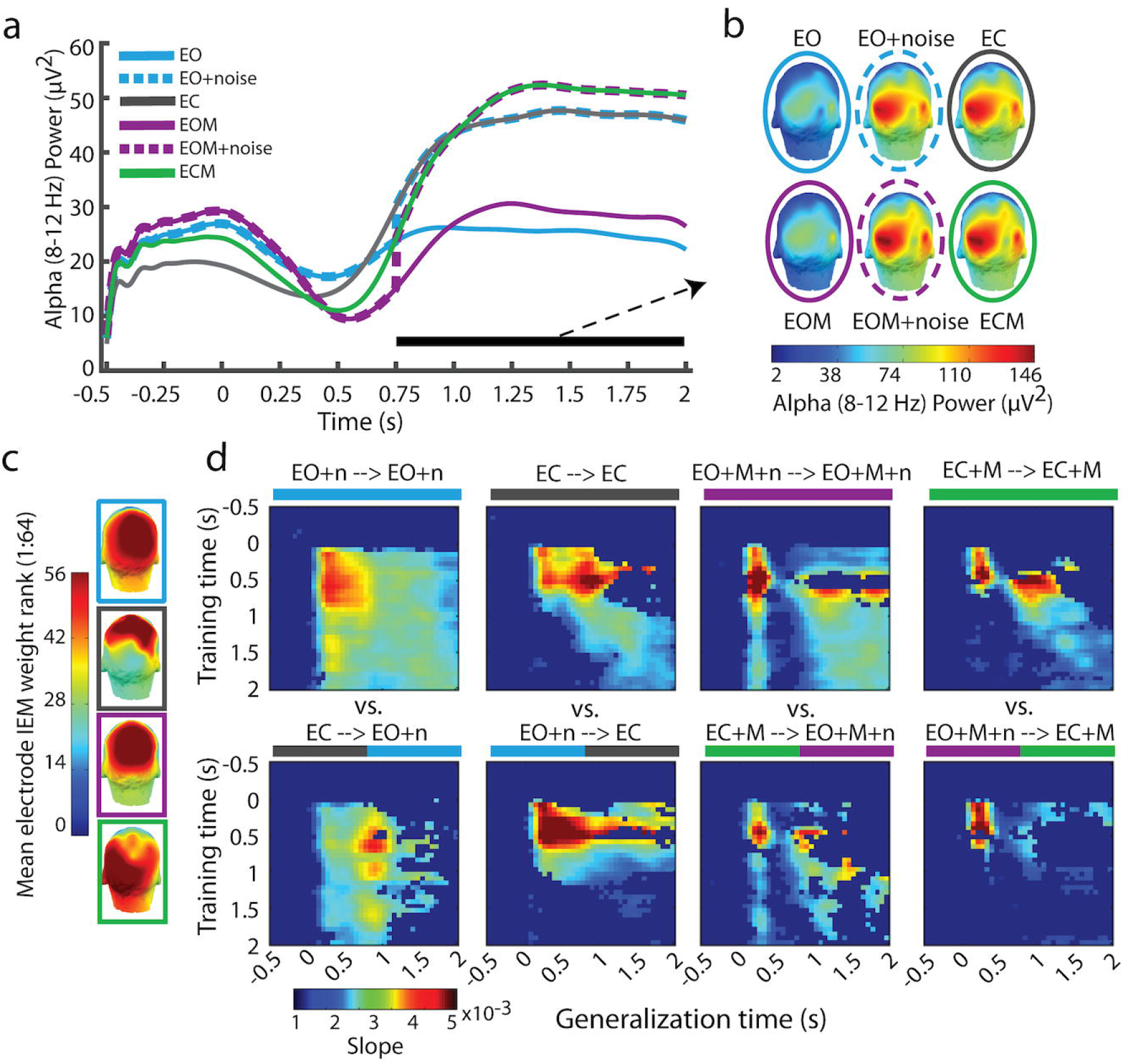
Investigation into the role of alpha enhancement with eye closure on temporal generalization within and between conditions. **(a)** Alpha power by condition demonstrating the alpha enhancement with eye closure and its recreation in the eyes open conditions for the simulation. **(b)** Scalp topography of alpha power by condition demonstrating that alpha is largely occipital in all conditions, and the simulation of alpha power topography in the eyes open conditions. **(c)** Scalp topography of electrode weight rank, where higher values indicate larger weights for those electrodes resulting from the IEM. **(d)** The results of the temporal generalization with the simulated recreation of alpha enhancement in the eyes open conditions. The pattern of temporal generalization is unchanged by this simulated alpha enhancement (see Fig. 3).

In order to examine this possibility, we again ran the IEM and the temporal generalization analysis for the eyes open conditions, but prior to modeling we added alpha power to simulate the increase in alpha power observed in the eyes closed condition. Specifically, in the eyes open conditions we added a random amount of power, to each electrode, at each time point from 750 (deadline for eye closure) to 2,000 ms after target onset. This random power was sampled from a normal distribution with the mean of the difference in power between the eyes closed and eyes open conditions, and the pooled standard deviation of power in the eyes open and eyes closed conditions *for that electrode at that time point*. This added noise reflects any systematic differences in alpha power (see **Fig. 5a**) and topography (see **Fig. 5b**) resulting from eye closure.

In other words, the added noise recreates the exact changes in alpha power and topography evoked by eye closure in the eyes open conditions. However, as the noise is drawn *randomly* from the distribution of difference in power and topography between eyes open and eyes closed conditions, this noise should not impact the temporal generalization of the spatially selective representation. This is necessary the case, as the spatially selective representation extracted by the IEM is based on the pattern of alpha that is predicted by the *a priori* defined set of spatially selective neural channels (the basis set). In order for *any* change to alpha power and/or topography to affect the results of the IEM, and its generalization, it must necessarily impact the relationship between the pattern of alpha power and the basis set (i.e., the beta weights). Changes that are independent of the pattern of alpha that codes the spatially selective representation will not affect the result of the IEM or its generalization.

Alpha power in the eyes open conditions with the added noise resembled that in the eyes closed condition (**Fig. 5a**). The topography of alpha power in the eyes open conditions with the additional noise also closely resembled that of the corresponding eyes closed condition during the 750-2,000 ms period (**Fig. 5b**). The results of the temporal generalization of the IEM in the eyes open conditions (both with and without mask) with the added noise were identical to those observed without (**Fig. 5d**). This simulation supports the conclusion that changes in alpha power and topography resulting from eye closure that are layered on top of the patterns of alpha oscillations coding remembered locations, but are otherwise independent of the memory activity, are insufficient to disrupt the IEM reconstructions of remembered locations and their generalization.

### Is process 2 evoked by eye closure or by the absence of continued visual input?

The purpose of the eyes closed conditions was to examine whether the absence of continued visual input would affect the encoding of spatial information in alpha-band activity. However, as discussed in the previous section, it is possible that the effects we observe are specific to some phenomenon unique to eye closure. We have shown that the well-documented effect of increased alpha power associated with eye closure does not account for the pattern of temporal generalization where process 1 and 2 are evident. As a further control we collected data from an additional independent sample from two conditions (**Fig. 6a**): (1) the same eyes open condition as in the main study, and (2) an eye-blink condition.

**Figure 6.**
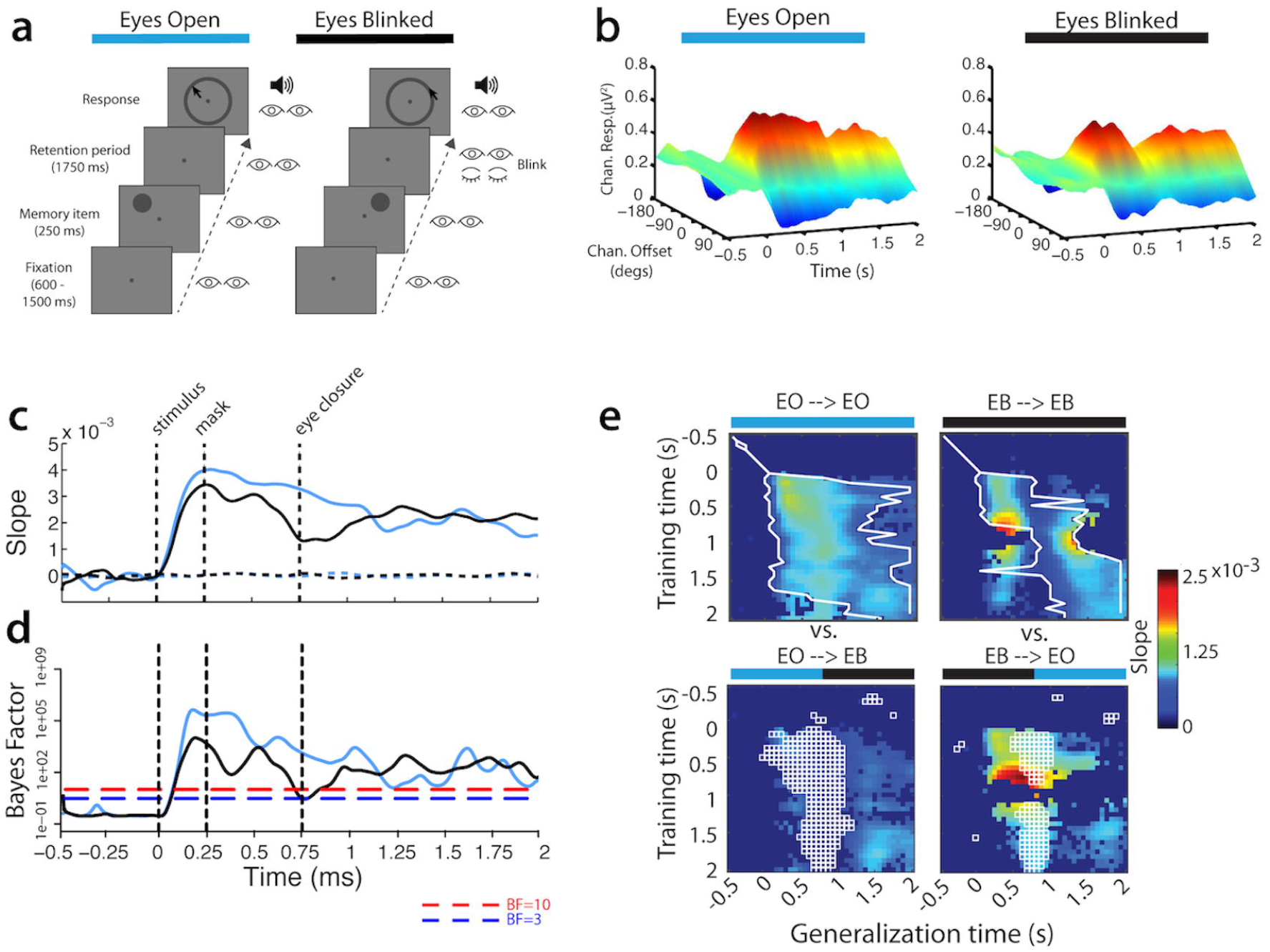
Control study into the effects of eye closure (blink). **(a)** Trial procedure for each experimental condition. **(b)** Estimated neural population response, or “channel” (offset of channel’s preferred location from stimulus location: ±0,45,90,135, or 180°), response (alpha-band power) starting 500 ms before onset of stimulus until end of retention period. **(c)** Linear regression weights of estimated channel response folded around 0° offset (“slope”), indicating the degree of spatial selectivity. **(d)** Bayes Factor (BF) of t-test comparing slope at each time point to slope of response estimated using permuted location labels. **(e)** Results of temporal generalization.

There was evidence of a spatially selective response, centered on the stimulus location, in the alpha band following stimulus presentation in both conditions (**Fig. 6b/c**). Interestingly, as in the eyes open masked condition, that spatially selective response was briefly disrupted by the eye-blink but recovered to a degree indistinguishable from the eyes open condition.

The evidence for a unitary process (process 1) was replicated in the eyes open condition, and, again as in the eyes open masked condition, that same process was evident throughout the eye-blink condition except for the brief disruption following the blink (**Fig. 6d**). Importantly, we did not observe the pattern that would have indicated the emergence of process 2 in the eye-blink condition, as we did in the eyes closed conditions of the main study. Thus, we conclude that the emergence of process 2 is not a result of eye closure, but rather the continued absence of visual input during the retention interval.

### What is different between process 1 and process 2?

In order to address this question we first examined whether there were any differences in the topography of alpha power as a function of condition and time window – i.e., either those times overlapping with process 1 (during cue presentation) or process 2 (retention interval). Given previously reported results we should expect that alpha power is greater at electrode sites ipsilateral to the cue location as compared to contralateral sites – indicating that spatial attention/memory modulated the topographical distribution of alpha (Kelly et al., 2006; Sauseng et al., 2005; Thut et al., 2006; Worden, Foxe, Wang, & Simpson, 2000). To that end we examined the alpha lateralization as a function of condition and time window.

There was significant alpha lateralization such that alpha power was greater at ipsilateral sites than at contralateral sites (effect of laterality, BF = 231.85; **Fig. 7a**). Alpha lateralization was greater for eyes open conditions than eyes closed conditions (interaction of eyes and laterality, BF = 18.64; **Fig. 7b**). The lateralization of alpha was also reduced during the time window overlapping with the emergence of process 2 relative to that overlapping with process 1 (interaction of time window and laterality, BF = 79.88). However, the effect of eyes did not modulate the effect of time window on laterality (all BFs < 1). Thus, there is no difference in the lateralization of alpha between eyes open and eyes closed conditions unique to the retention period, which could have possibly account for the difference between process 1 and 2.

**Figure 7.**
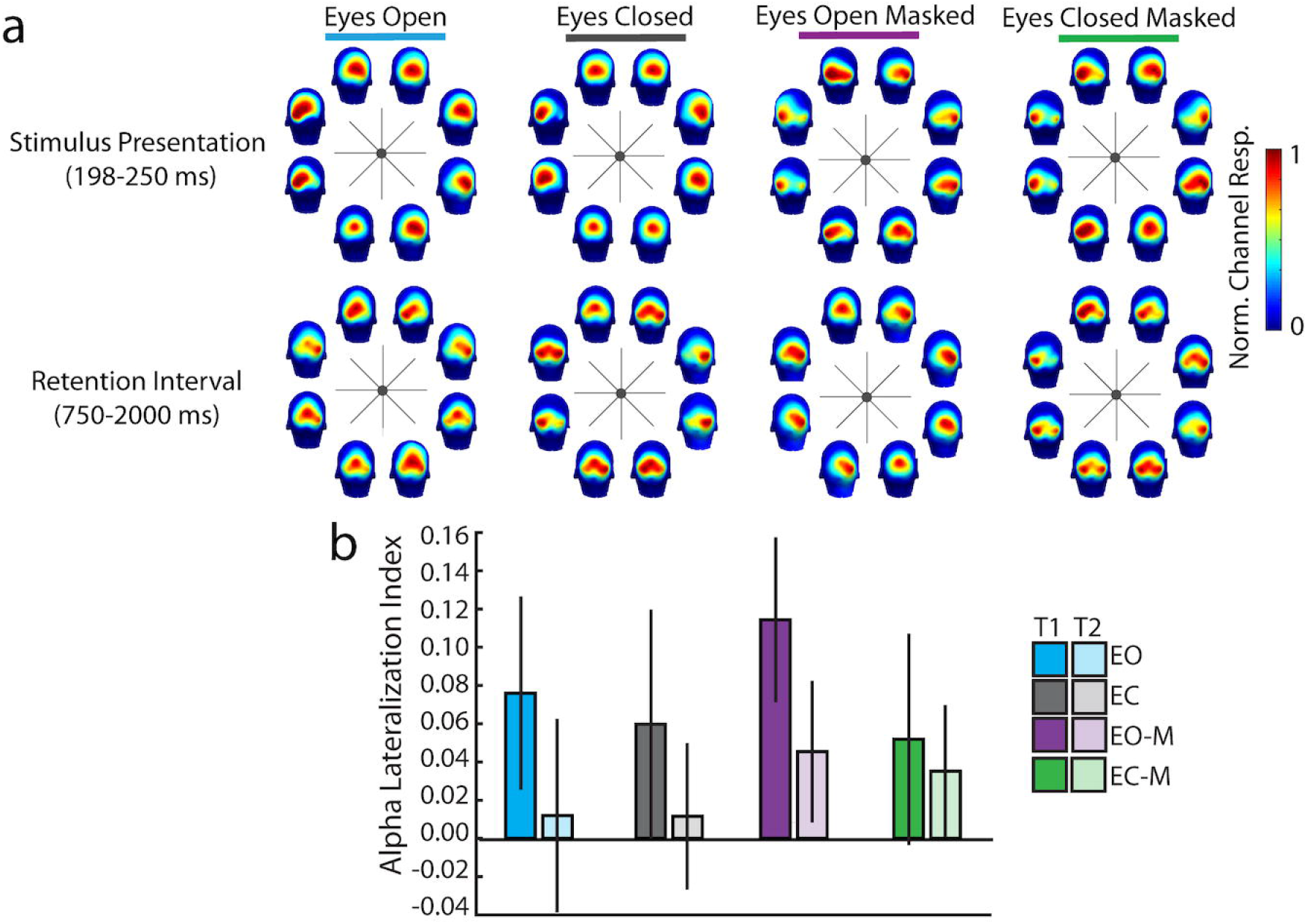
(a) Topography of normed alpha power during process 1 and process 2 time windows as a function of spatial location and condition. Alpha power is normalized to the range of alpha at P-O electrode sites (i.e., proportion of max value of alpha power among P-O electrodes for that condition) in order to be able to observe local topography for those critical sites. (b) Alpha lateralization index (alpha power at ipsilateral sites – contralateral sites / ipsilateral + contralateral) by condition and time window - overlapping with process 1 (T1 = 198-250 ms) or process 2 (T2 = randomly selected 52 ms window from 750-2,000 ms).

Within the context of the analytical approach we employed here (IEM), the difference between process 1 and process 2 is that the pattern of GLM weights across electrodes generated by the IEM differs between process 1 and process 2 in the eyes closed conditions, but not within process 1 and process 2. Indeed in the eyes open condition there is no consistent change in the weights beginning at ∼300 ms after target onset – corresponding to process 1 (**Fig. 8a**). In contrast, in the eyes closed condition there are two separate time windows during which there is no consistent change in weights: the first from ∼300 to 900 ms after target onset, and the other from ∼900 to 2,000 ms. These two time periods roughly correspond to the two processes evident in the temporal generalization of the IEM. From the first time window, possibly equating to process 1, to the second time window, possibly equating to process 2, weights on average moved in the negative direction.

**Figure 8.**
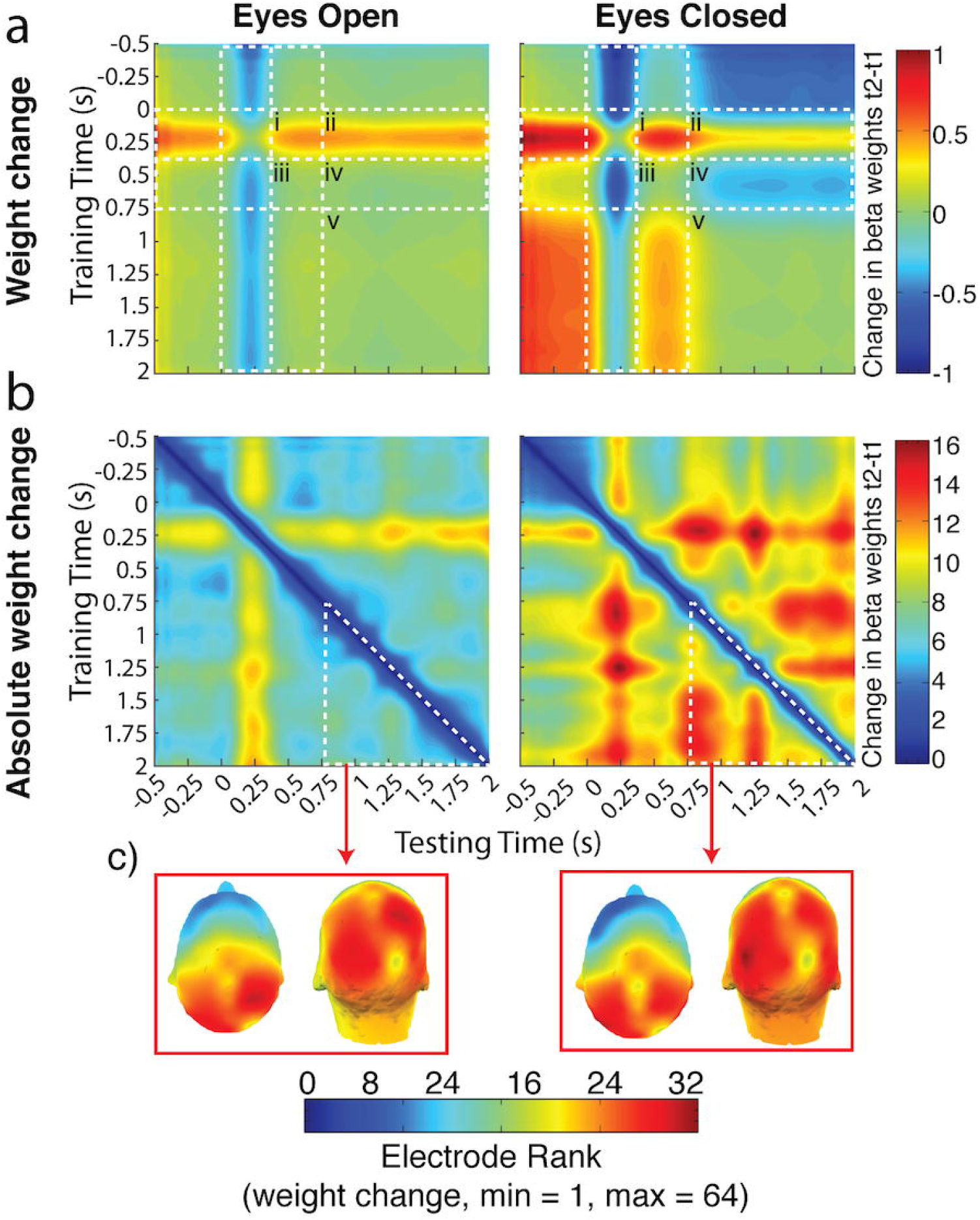
Investigation into the nature of the weight change. (a) In the **eyes open** condition average weight change was ∼0 throughout the retention interval (*iii-v*), following the initial average increase in weights from 250 ms after stimulus onset forward in time (*i* and *ii*). This same increase in weights was also present in the **eyes closed** condition (*i* and *ii*). However, there was an additional average change unique to the **eyes closed** condition - a decrease from around the time of eye closure (∼300 to 750 ms after stimulus onset) forward into the rest of the retention interval (*iv*). Otherwise average weight change in the **eyes closed** condition was ∼0 (*iii* and *v*). (b) Weights were more dynamic throughout the trial in the **eyes closed** condition, including prior to eye closure, than in the **eyes open** condition. (c) The changes in the **eyes closed** condition after eye closure, while greater than that during the same time period in the **eyes open** condition, are largely in parietal and occipital electrodes as they are in the **eyes open** condition.

Weights were more dynamic, showing greater absolute change over time, in the eyes closed conditions (**Fig. 8b**), especially during the retention interval, than in the eyes open conditions. In order to examine the topography of the change in weights over time during the retention interval we ranked the electrodes according to absolute change in weight between the two time points of every sample pair in the temporal generalization matrix. We then averaged these rankings over the generalization matrix during the period from 750-2,000 ms. For both eyes open and eyes closed conditions parietal and occipital electrode sites were consistently ranked highest for weight change (**Fig. 8c**). These electrodes also consistently rank highest in terms of absolute weight in the IEM (**Fig. 8d**). This suggests that the differences in weights between process 1 and process 2 are more likely to be located at parietal and occipital electrode sites, although the exact nature and meaning of these weight differences is unknown.

## Discussion

Alpha oscillations are the predominant systematic and recurring voltage fluctuation in the human electroencephalogram and were first described by Berger (1930). While alpha oscillations were initially considered to represent a low arousal state (e.g., ‘cortical idling’, Adrian & Matthews, 1934; see also Pfurtscheller, Stancák, & Neuper, 1996), it is now clear that oscillations in the 8-14 Hz range play a key role in perception and cognition, perhaps via mechanism of inhibition (Jensen & Mazaheri, 2010; Klimesch, 2012), although perhaps not (see Foster & Awh, 2019). Selective attention to, and retention of, behaviorally relevant locations can both be measured via alpha oscillations (Foster et al., 2016, 2017; Kelly et al., 2006; Medendorp et al., 2007; Samaha et al., 2016; Sauseng et al., 2005; Thut et al., 2006; van Moorselaar et al., 2018; Worden et al., 2000), suggesting that alpha oscillations represent the functioning of a unitary, or at the very least a shared, mechanism for spatial representation involved in both attention and memory. The evidence presented here, regardless of analytic approach (encoding - IEM or decoding – linear discriminant classification), however, suggests that alpha does not support a single unitary cognitive mechanism, but rather that there are at least two distinct processes that support the selection and retention of behaviorally relevant locations that operate within the alpha frequency band: one that is fast and continuous, supporting selection of information from the sensory environment and that is disrupted by masking and no sustained visual input; and a second that is delayed relative to the first, emerging upon the cessation of sustained visual input.

### Alternatives

While the present results are consistent with the notion that alpha supports at least two distinct processes supporting the mental representation of behaviorally relevant locations, facets of the experimental design introduce known artifacts that may appear as confounds. First, the mere closing of the eyes results in a robust and reproducible increase in alpha power that varies systematically across the scalp, being larger at posterior electrodes than at other electrodes (see **Fig. 5a**). These dramatic modulations in alpha oscillations, the signal encoding spatially selective representations, are an obvious candidate for the disruption to inter-temporal generalization. Importantly, when alpha power and topography in the eyes open condition was made to resemble that of the eyes closed condition during the retention interval as noise, the results of the temporal generalization are unchanged (**Fig. 5b**). This simulation demonstrates that changes in alpha power and topography, such as those that follow eye closure, cannot influence the results of the IEM, or its temporal generalization, when uncorrelated with the basis set (i.e., unrelated to the pattern which encodes location). Thus, it is unlikely that the lack of temporal generalization reflects a contamination of alpha oscillations that actually represent a unitary mechanism, rather we argue that the pattern of temporal generalization (and lack thereof) is evidence for at least two alpha-based processes involved in the selection and retention of spatial information.

Eye closure also results in large ocular artifacts that propagate across the scalp. Thus, it is reasonable to suggest that the impaired generalization is due to residual artifacts due to eye closure. However, we demonstrated in an independent control study that mere eye closure (as when blinking), and any artifacts associated with that ocular event, do not yield the same pattern of temporal generalization evident in the eyes closed conditions (**Fig. 6**); rather, the temporal generalization matrix in the eye-blink condition resembled that of the eyes open-masked condition, i.e., a single process temporarily disrupted.

The results of the eye-blink control study also address the possibility that the difference between the eyes open and closed conditions is attributable to a dual-task demand of having to close their eyes, i.e., as reconstructions of remembered spatial locations are sensitive to demands on attention (van Moorselaar et al., 2018). While it is true that participants had to close their eyes and wait for the tone to give the response, participants experienced a similar level of demand in the eye-blink condition. Furthermore, all conditions required participants to hold the information in memory and wait until the tone indicated it was time to give the response, and even the eyes open conditions made an additional demand (maintain fixation). Thus, the conditions are quite closely matched in terms of divided attention demands, making this account no more, and perhaps less likely than the guaranteed effect that continuous external visual input was absent in one condition and not the other.

This alternative would also predict that the reconstructed profiles of spatially-selective representations in those conditions with increased demands on attention (i.e., eyes closed), would be degraded relative to conditions without the divided attention demands (i.e., eyes closed). However, the results, as depicted in **Fig. 2**, reveal that the slopes of the reconstructed spatially-selective channel responses in the eyes open and eyes closed conditions do not differ during that period when process 2 emerges. Thus an account based on differential divided attention demands is not a viable alternative explanation of our findings.

In addition to the issue of noise, and related to the issue of reduced SNR, losses in temporal generalizability could also be attributed to unobserved losses in the signal of interest following eye closure. However, the pattern of temporal generalization in the eyes closed condition, in this case, cannot be explained by signal loss. The within condition IEMs during this time window reliably reconstruct the spatial location held in memory in the eyes closed condition to the same degree as in the eyes open condition. Thus the signal that carries the location information is present to the same extent in the eyes closed condition as in the eyes open condition during the same period of the retention interval when the loss of temporal generalization occurs.

## Conclusion

Based on the collection of evidence presented here we suggest that mental representations of stimulus location are retained via at least two distinct processes coded in alpha band oscillations. When visual input continues to be available, spatially selective representations can be supported and even recovered by the same process present when a location is initially selected and may be mediated by “attention-based rehearsal” (Awh, Jonides, & Reuter-Lorenz, 1998; Awh et al., 1999; Postle, Awh, Jonides, Smith, & D’Esposito, 2004) of the *external* sensory environment, as has previously been observed (Foster et al., 2016, 2017; van Moorselaar et al., 2018). However, when visual input is not available, a process possibly based on *internal* selection from information encoded in memory (Chun, Golomb, & Turk-Browne, 2011; Kiyonaga & Egner, 2013) emerges to support the ongoing spatial representation. To the extent that the topographic pattern of alpha oscillations is reflective of the activity in retinotopically mapped brain regions (de Munck et al., 2007; Kelly et al., 2006; Worden et al., 2000), then our results suggest that these areas are differentially activated when eyes are closed than when eyes are open. This differential involvement may be the result of local changes in patterns of activity (e.g., local patterns of inhibitory activity) or from perturbations from other brain areas. The latter possibility would be consistent with recent evidence that initial selection of remembered information is supported by oscillatory activity in prefrontal and parietal/occipital locations, while the reactivation of that information in memory relies a network of occipital/temporal locations (Quentin et al., 2019).

## Acknowledgements

This work was supported by the Institute for Collaborative Biotechnologies through contract W911NF-09-0001 and cooperative agreement W911NF-19-0026, both from the U.S. Army Research Office. The content of the information does not necessarily reflect the position or the policy of the Government, and no official endorsement should be inferred. We would like to thank James Elliott for his crucial feedback and suggestions, as well as Caroline Trinh, Lena Nalbandian, and Scott Patterson for their help with data collection. M.H.M. and T.B. are co-first authors. M.H.M, T.B., and B.G. designed the experiment. T.B. programmed the experiment. M.H.M and T.B. collected the data. T.B., M.H.M., and B.G. analyzed the data. M.H.M, T.B., and B.G. wrote the manuscript.

